# Interferon signaling underlies radiotherapy responses in malignant peripheral nerve sheath tumors (MPNSTs)

**DOI:** 10.1101/2025.03.09.642167

**Authors:** Iowis Zhu, Julian Chien, Kanish Mirchia, Sixuan Pan, Kaeli Miller, Joanna Pak, Rosanna Wustrack, Varun Monga, Steve E. Braunstein, Line Jacques, Melike Pekmezci, S. John Liu, Harish N. Vasudevan

## Abstract

Malignant peripheral nerve sheath tumors (MPNSTs) have poor outcomes despite multimodal treatment with surgery, radiation, and systemic therapy. Here, we combine bulk and single cell transcriptomics, genome wide CRISPRi screens, and multiplatform molecular analysis across MPNST cells, mouse allograft models, and human patient samples to understand the mediators of radiation response. Our data reveal MPNSTs induce a type I interferon signature that functionally underlies radiation response. Moreover, irradiation of immunocompetent mouse MPNST allografts leads to interferon mediated T-cell recruitment and activation. Analysis of human MPNST resection specimens demonstrates increased microenvironmental and CD8+ T-cell infiltration are associated with improved local control following radiotherapy while chromosome 9p loss, which harbors an interferon gene cluster, is associated with poor radiotherapy response and decreased CD8+ T-cell infiltration. Taken together, these results provide preclinical rationale for combining immunomodulatory agents targeting interferon signaling to improve radiation responses in MPNSTs, which may be broadly applicable to other soft tissue sarcomas.

## Main Text

Neurofibromatosis type 1 (NF-1) is an inherited condition affecting approximately 1 in 3000 individuals worldwide.^1^ People with NF-1 are at increased risk for developing tumors of the peripheral nervous system (PNS) including benign plexiform neurofibromas (pNFs) that can undergo transformation into malignant peripheral nerve sheath tumors (MPNSTs). MPNSTs are the most common cause of death in adults with NF-1.^2^ Despite multimodal therapy with surgery, radiation and systemic therapy, outcomes remain poor for people with MPNSTs,^3,4^ thus comprising an urgent, unmet clinical need. In addition, people with NF-1 are at increased risk for secondary malignancies following radiotherapy,^5^ motivating an improved understanding of the therapeutic window for radiotherapy in this patient population.

MPNSTs are clinically managed as soft tissue sarcomas, a heterogeneous class of tumors treated with surgery, radiation, and systemic therapy, based on patient and tumor specific factors.^6,7^ Given the central role of radiotherapy in the management of sarcomas, multiple groups have developed nomograms predictive for radiation response that integrate clinical or pathologic factors,^8,9^ including specifically for MPNSTs.^3,10^ However, these frameworks do not account for genetic mutations or microenvironmental composition. Indeed, combining the immune checkpoint inhibitor pembrolizumab with preoperative radiation therapy improves outcomes compared to radiotherapy alone for undifferentiated pleomorphic sarcomas or liposarcomas,^11^ underscoring the importance of the immune microenvironment in sarcoma treatment. However, whether this applies to other sarcoma subtypes such as MPNSTs is not well understood. More broadly, the functional mediators and genetic basis for differential radiation response, as well as the mechanisms underlying the synergy between radiotherapy and immunomodulatory treatment approaches, remain unclear.

In this study, we sought to elucidate mechanisms underlaying radiation response in MPNSTs by combining bulk RNA-sequencing, genome-wide CRISPR interference (CRISPRi) screens, single cell RNA-sequencing (scRNA-seq) of immunocompetent murine MPNST allograft models, and genomic analysis of human MPNST specimens. We found that MPNSTs are radioresistant and induce a distinct transcriptional signature compared to benign pNFs, as highlighted by enrichment for immunomodulatory pathways such as interferon signaling. Consistent with this observation, genome wide CRISPRi screens converged on type I interferon (IFN) signaling as a critical mediator of radiation response. ScRNA-seq of irradiated murine MPNST tumors *in vivo* demonstrated that radiation increased T cell recruitment and activation with concomitant induction of interferon secretion both by tumor cells and by macrophages within the microenvironment. Finally, analysis of human MPNST specimens from patients treated with surgery and radiation revealed that radiation response was associated with increased microenvironmental infiltration, CD8+ T-cell presence, and chromosome 9p loss, which harbors an interferon gene cluster. Taken together, our study demonstrates the importance of interferon signaling from both tumor cells and the microenvironment in mediating radiation responses in MPNSTs, providing preclinical rationale for stimulating this interferon response as an approach to improve the efficacy of radiotherapy for sarcomas.

### MPNST cells demonstrate increased radioresistance and modulate interferon signaling pathways

To measure the single fraction radiation dose response in pNF and MPNST, two human MPNST cell lines, ST88-14^12^ and JH-2-002,^13,14^ and the human pNF cell line NF9511.b^15^ were treated with a single radiation dose ranging from 0-8 Gray (Gy) and stained with the viability dye DRAQ7 for flow cytometry (Figure 1a). MPNST cells demonstrated significantly increased viability at 4 Gy and 8 Gy compared to pNF cells. To measure survival following fractionated radiotherapy, cell lines were similarly treated with daily treatment of 2 Gy for up to 5 total days and stained with the viability dye DRAQ7, which demonstrated MPNST cells had significantly increased viability across multiple timepoints (Figure 1b). Taken together, these data demonstrate MPNST cells are radioresistant compared to pNF cells.

**Figure 1.**
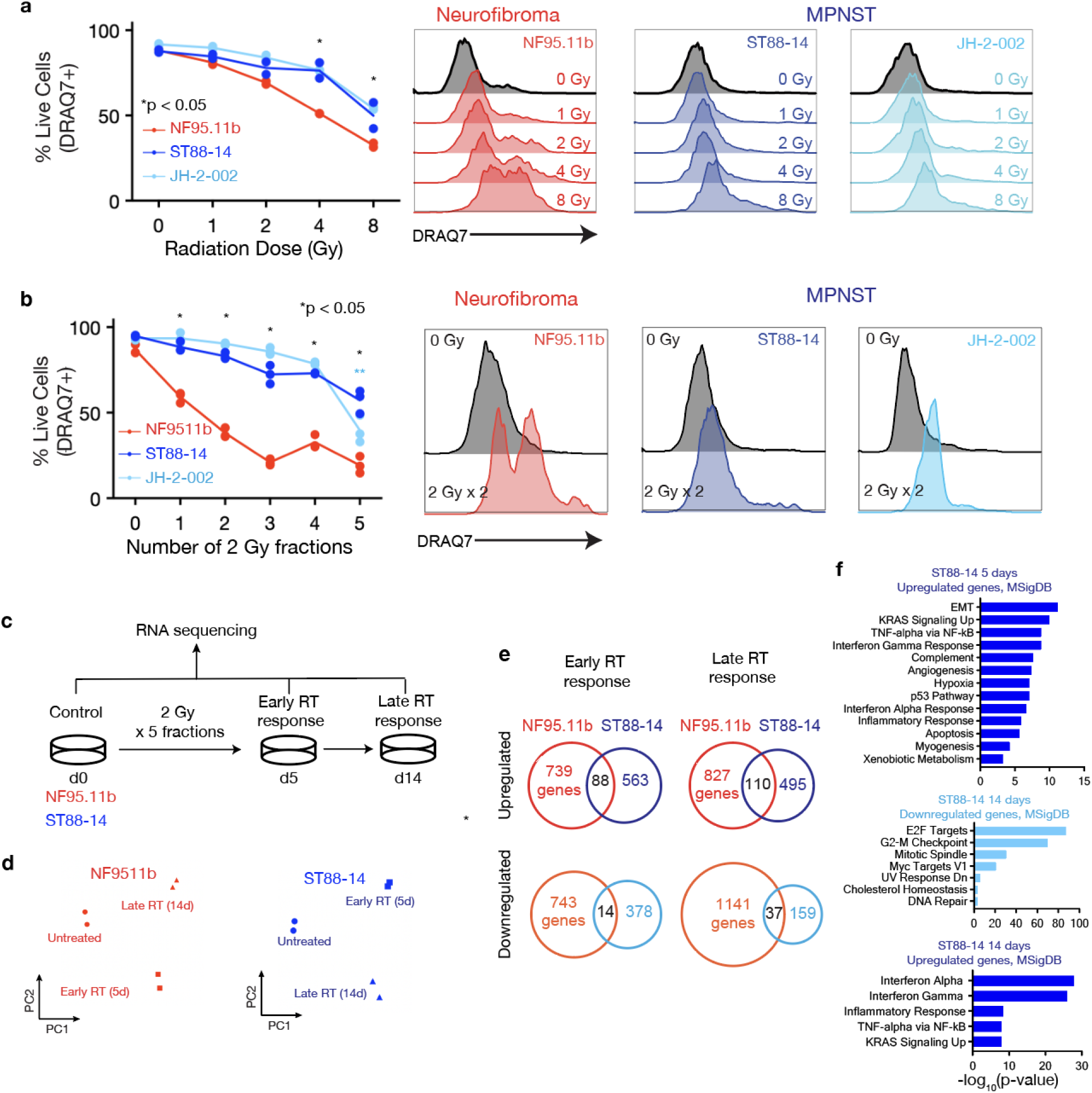
MPNST cells demonstrate increased radioresistance compared to pNF cells and transcriptionally modulate interferon signaling. a. Left: Percentage of live cells as measured by DRAQ7 staining for NF95.11b pNF cells compared to ST88-14 and JH-2-002 MPNST cells treated with single dose irradiation at Day 0 and harvested at Day 4. Right: Representative histograms of DRAQ7 staining of the indicated cell lines at different radiation doses as compared to Day 0. b. Left: Percentage of live cells as measured by DRAQ7 staining for pNF and MPNST cell lines treated with the indicated number of daily 2 Gy fractions and harvested 48 hours after completion of irradiation. Right: Representative histograms of DRAQ7 staining of the indicated cell lines after 2 fractions of 2 Gy. c. NF95.11b pNF and ST88-14 MPNST cell lines were irradiated with 5 fractions of 2 Gy and harvested at 5 days and 14 days post irradiation for bulk RNA sequencing. d. Principal component analysis of bulk RNA sequencing data reveals radiation modulates the transcriptome in both NF95.11b pNF and ST88-14 MPNST cells. e. Comparison of significantly upregulated and downregulated genes between cell lines and timepoints shows unique gene expression changes in NF95.11b pNF compared to ST88-14 MPNST cells. f. Gene ontology (GO) analysis of differentially regulated genes in ST88-14 MPNST cells during early RT response (5 days) and late RT response (14 days) reveals induction of interferon signaling and repression of cell cycle progression. * p<0.05 by Student’s t test.

To identify transcriptomic changes between pNF and MPNST cell lines post-RT that underlie the observed differences in radiotherapy response, we next irradiated the pNF cell line NF9511.b and the MPNST cell line ST88-14 with 5 daily fractions of 2 Gy and collected cells at day 5 and day 14 post-RT for bulk RNA-sequencing (Figure 1c; Supplementary Table 1). Principal component and differential gene expression analysis of RNA-sequencing data revealed radiation globally modulated the transcriptome in both cell lines (Figure 1d; Supplementary Figure 1). Distinct sets of significantly upregulated and downregulated genes were observed in response to radiation in pNF and MPNST cells (Figure 1e). Using gene ontology (GO) analysis of these significantly regulated gene sets, we found that ST88-14 MPNST cells, but not NF9511.b pNF cells, significantly upregulated IFN signaling genes during both early and late RT response and downregulated mitosis-associated genes in response to RT (Figure 1f). In sum, MPNST cells demonstrate radioresistance compared to pNF cells and exhibit a distinct transcriptomic response to radiation enriched for IFN signaling transcriptional signatures.

### Genome wide CRISPRi screen reveals contributors to MPNST radiotherapy response

While our RNA-sequencing revealed changes in IFN gene signatures associated with MPNST radiation response, it remained unclear whether these differences were functionally required for radiation responses in MPNST cells. Thus, we next performed a genome wide CRISPRi screen to identify functional mediators of radiation response in JH-2-002 MPNST cells. To individually knock down genes in an unbiased fashion, we transduced a barcoded lentiviral dual single guide RNA (sgRNA) library into JH-2-002 human MPNST cells expressing dCas9-Zim3, with virus titrated to approximately 1 sgRNA vector per cell. Cells were sorted for library expression using FACS (Supplementary Figure 2a), and a T0 control was collected at Day 0. Two experimental groups were designed, with one receiving 2 daily fractions of 2 Gy and the other being untreated, receiving no radiation. Genomic DNA (gDNA) from all cells was harvested on Day 14 and sgRNA barcodes were amplified and sequenced (Figure 2a; Supplementary Table 2). To identify essential genes required for JH-2-002 MPNST cell growth, we first compared untreated T14 sgRNA abundances to T0 sgRNA abundances (Untreated T14 / T0), identifying 985 significantly enriched sgRNAs mediating increased cell growth and 323 significantly depleted sgRNAs mediating decreased cell growth (Supplementary Figure 2b). SgRNAs that inhibited cell growth were enriched for DNA repair pathways and chromatin remodeling (Supplementary Figure 2c) with noted genes *RREB1, PARP1*, and *KAT2B* (Supplementary Figure 2d). As expected, sg*TP53* repression significantly promoted cell growth (Supplementary Figure 2e).

**Figure 2.**
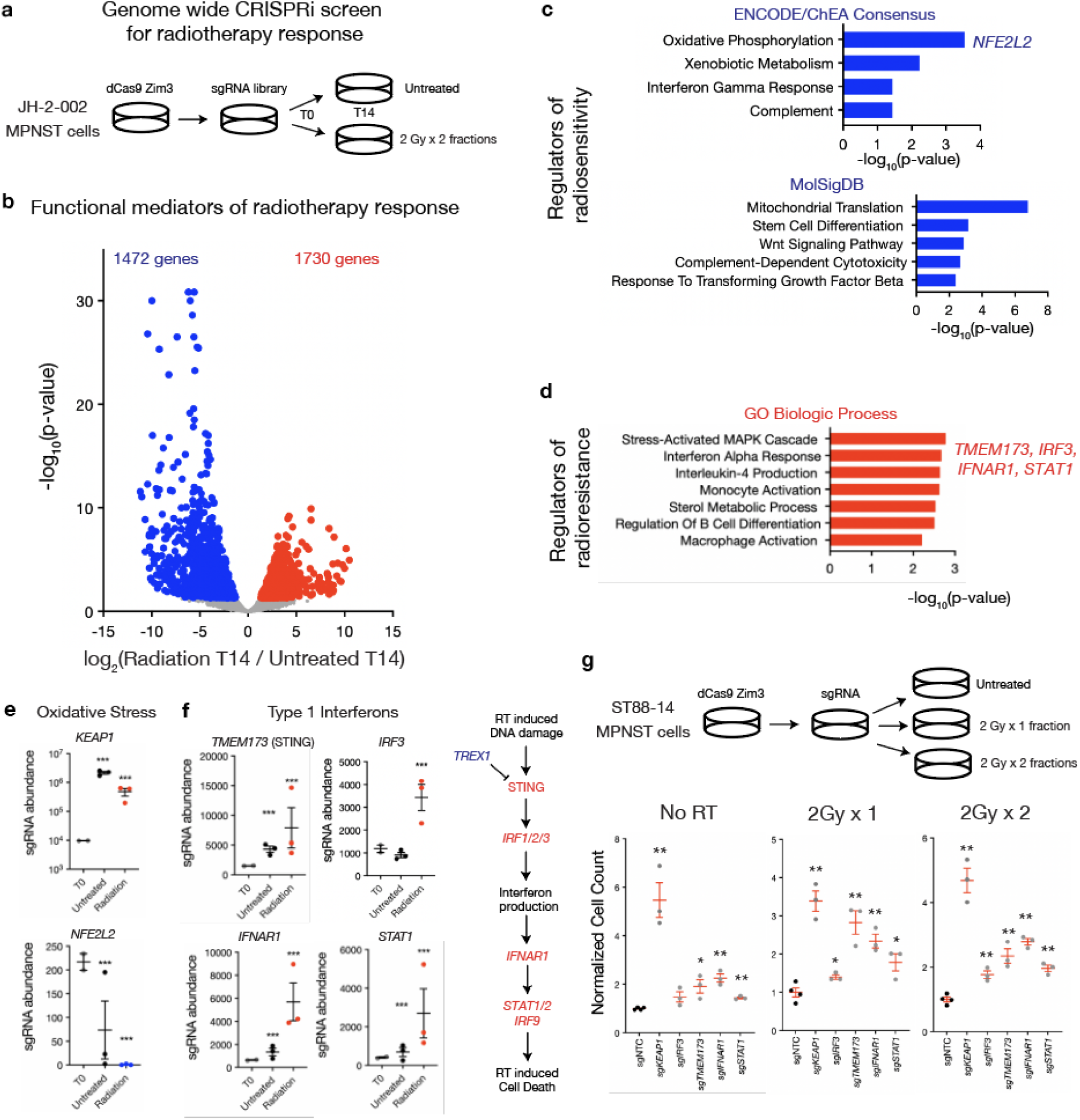
Genome wide CRISPRi screen reveals type 1 interferons underlie radioresistance in MPNST cells. a. Experimental setup of CRISPRi screen in JH-2-002 MPNST cells. b. Volcano plot depicting significantly enriched sgRNAs (n = 1730, red) or depleted sgRNAs (n = 1472, blue) between irradiated and control groups, p < 0.05. c. GO analysis of significantly depleted sgRNAs mediating radiosensitivity reveals enrichment for oxidative phosphorylation pathways (*NFE2L2)*. d. GO analysis of significantly enriched sgRNAs mediating radioresistance reveals enrichment for interferon signaling as well as multiple cytokine and immunomodulatory pathways. e. Normalized sgRNA abundances of *KEAP1* and *NFE2L2*, two members of the oxidative stress response. f. Normalized sgRNA abundances of members of the type I interferon signaling pathway: *TMEM173* (STING), *IRF3, IFNAR1*, and *STAT1*. g. Radiation responses in *sgKEAP1, sgIRF3, sgTMEM173, sgIFNAR1,* and *sgSTAT1* validate these genes as mediators of radiation response in human ST88-14 MPNST cell lines treated with 2 Gy for 1 or 2 fractions and harvested at 48 hours following completion of irradiation.* p<0.05 compared to sgNTC; ** p<0.01 compared to sgNTC, Student’s t-test.

A total of 1,730 sgRNAs were significantly increased in abundance after radiation (Radiation T14 / Untreated T14), suggesting these sgRNAs mediated radiation resistance, while 1,472 sgRNAs were significantly decreased in abundance, suggesting these sgRNAs mediated radiation sensitivity (Figure 2b). GO analysis revealed that oxidative phosphorylation and stress genes, including *NFE2L2* encoding the master oxidative stress response regulator NRF2, were significantly enriched amongst depleted sgRNAs mediating radiosensitivity while *KEAP1,* a negative regulator of NRF2, was significantly enriched following radiation (Figure 2c and 2e), consistent with the role of the Nrf2 signaling axis in modulating radiation response.^16,17^ Amongst genes significantly enriched in irradiated cells, components of the type I IFN response pathway, such as *TMEM173* (encoding the STING protein), *IRF3*, *IFNAR1*, and *STAT1*, were significantly enriched sgRNAs underlying radiation resistance (Figure 2d and 2f; Supplementary Figure 3a). In addition, sgRNAs encoding previously reported mediators of radiosensitivity such as DNA repair genes (*ATM, RAD51AP1*) and TGFβ *(TGFB1, SMAD4)* were significantly depleted following radiotherapy (Supplementary Figure 3b), supporting the overall robustness of our screen. To validate our findings in an orthogonal model, we tested selected CRISPRi screen radiation modifier genes (*KEAP1, IRF3, TMEM173, IFNAR1, STAT1*) in a second human ST88-14 MPNST cell line (Figure 2g; Supplementary Figure 3c). Thus, our CRISPRi screen data identified multiple functional mechanisms underlying radiation response in MPNSTs, including a key functional role for interferon signaling consistent with our bulk RNA-sequencing data.

### MPNST tumors demonstrate increased interferon and T-cell signaling signatures in response to RT in vivo

Given interferon signaling is critical for mediating crosstalk between the tumor and immune microenvironment,^18^ we next sought to better understand MPNST radiation responses in an immunocompetent *in vivo* model. We thus subcutaneously implanted wildtype C57/B6 mice with two different *Nf1/Tp53* double knockout murine MPNST cell lines, JW18.2 and JW23.3,^19–21^ and randomized them into two experimental groups: 5 daily treatments of 2 Gy versus untreated tumors (Figure 3a). Tumors receiving RT had significantly reduced growth (Figure 3b). To define the transcriptomic changes and cellular subpopulations mediating radiation response in these immunocompetent MPNST models, we next performed single cell RNA-sequencing (scRNA-seq) on tumors from mice implanted with JW23.3 MPNST cells treated with radiation (n=4) versus untreated (n=3) tumors at day 21. Uniform manifold approximation and projection (UMAP) analysis revealed 7 tumor clusters and 9 non-tumor clusters (Figure 3c; Supplementary Figure 4a-b; Supplementary Table 3) which were defined using cluster marker genes (Supplementary Figure 4c), automated cell-type classification (Supplementary Figure 4d),^22^ *Xist* expression restricted to host female mice rather than male tumor cells (Supplementary Figure 4e) and cell-cycle phase assignment (Supplementary Figure 4f). Of these clusters, only cluster 6, corresponding to T and NK cells, with expression of *PTPRC, CD4,* and *CD8* (Supplementary Figure 5a), was found to be significantly increased in irradiated tumors compared to unirradiated control tumors (Figure 3d). Flow cytometry analysis of the CD3+ T-cell subset in an independent experiment was notable for a significant increase in CD4+ T-cells and PD-1 expression following RT (Figure 3e), supporting the changes observed in our scRNA-seq data. Differential gene expression analysis of irradiated Cluster 6 T/NK cells compared to un-irradiated control Cluster 6 T/NK cells revealed enrichment of genes important for TCR signaling and T-cell activation such as *Gzmb* and *Zap70,* as well as type I IFN signaling such as *Irf7* and *Stat1* (Figure 3f, Supplementary Figure 5b). We next performed differential gene expression analysis across the macrophage cell clusters 0, 3, and 10 between irradiated and control tumors (Supplementary Figure 5c-d). GO analysis revealed a significant upregulation of type I and II IFN response pathways as well as a significant shift toward M1 macrophage polarization in irradiated macrophages compared to control macrophages (Figure 3g; Supplementary Figure 5e). Finally, we performed differential expression analysis across MPNST cell clusters to determine tumor cell autonomous mechanisms of radiation response. GO analysis of tumor cell clusters 1 and 4 were notable for significantly upregulated expression of NFE2L2 target genes, suggesting one mechanism of tumor cell intrinsic radioresistance occurs through an NRF2 mediated oxidative stress response consistent with our CRISPRi screen data (Figure 3h; Supplementary Figure 6a-b). Finally, irradiated Cluster 7 tumor cells were enriched for genes encoding IL-2 and the type I IFN response pathway, supporting the importance of tumor cell autonomous interferon signaling in mediating radiotherapy responses as observed in our bulk RNA-sequencing and CRISPRi screen data (Figure 3h, Supplementary Figure 6c). No significant differentially regulated gene sets between irradiated and control cells were observed in the remaining tumor cell clusters. In sum, scRNA-seq of irradiated MPNST immunocompetent allografts reveals both tumor intrinsic and microenvironmental type I IFN secretion underlying T cell recruitment following radiotherapy.

**Figure 3.**
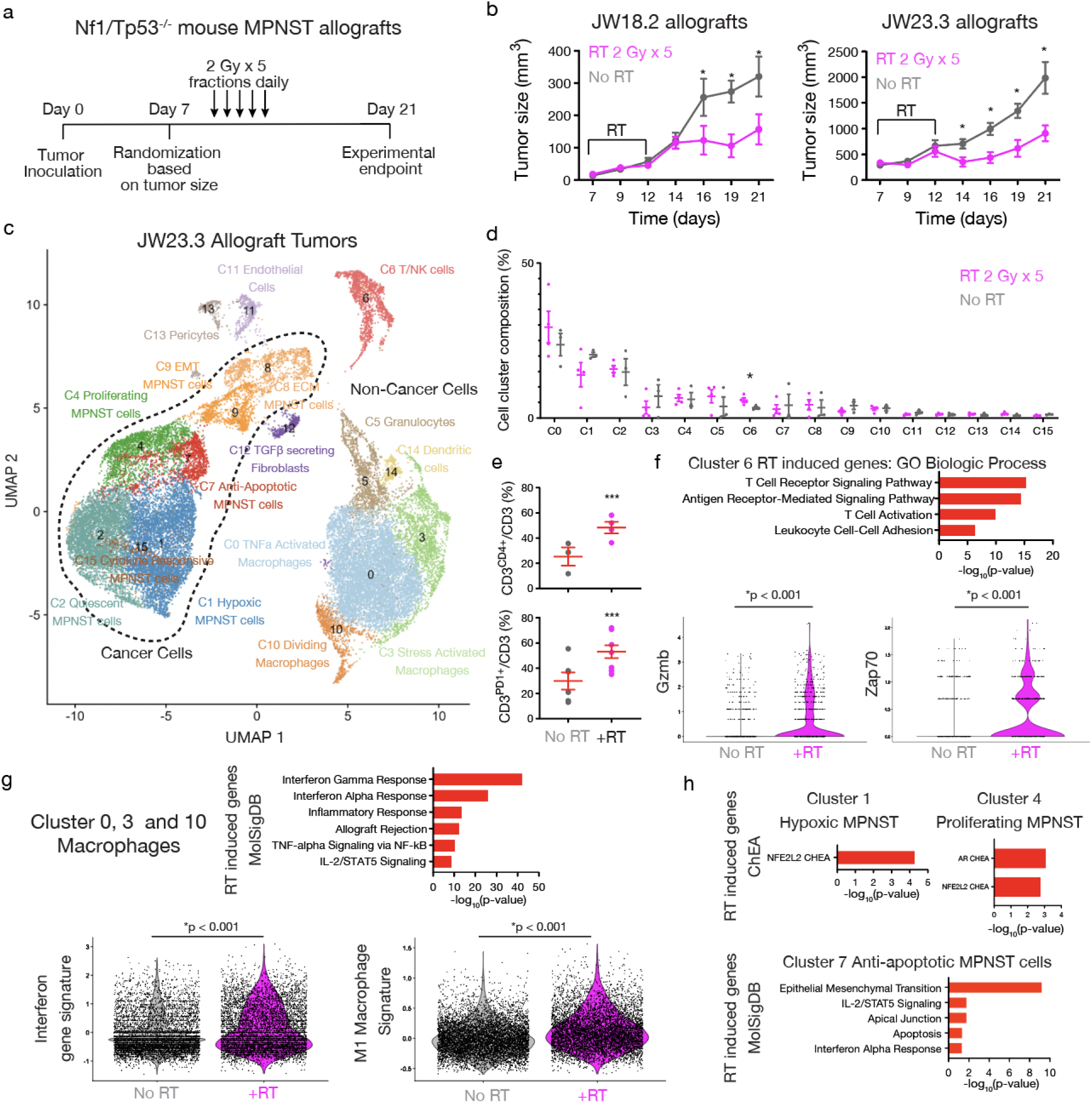
Irradiated mouse MPNST tumors demonstrate increased interferon activation and T-cell recruitment *in vivo*. a. MPNST tumor irradiation *in vivo* experimental design with mouse *Nf1/Tp53* mutant MPNST cells implanted subcutaneously in a C57/B6 wildtype mouse. b. Growth curves of implanted JW18.2 and JW23.3 murine MPNST tumors reveal decreased tumor volume following irradiation (*p<0.05, Student’s t test). c. Harmonized single-cell RNA sequencing uniform manifold approximation and projection (UMAP) of tumor and non-tumor cell clustering of 32,763 cells harvested from JW23.3 tumors (n=3 control, n=4 irradiated) identifies 7 tumor and 9 non-tumor clusters. d. Breakdown of cell cluster composition by experimental groups reveals only Cluster 6 T/NK cells are significantly enriched in irradiated tumors (*p<0.05, Student’s t-test). e. Flow cytometry analysis of top: Percentage of CD3+ cells that were also CD4+ between experimental group and bottom: Percentage of CD3+ cells that were also PD1+ between experimental groups reveals enrichment of both populations in irradiated tumors (*** p<0.01, Student’s t-test). f. Differential gene expression analysis of irradiated versus control Cluster 6 T/NK cells followed by Top: GO biological process analysis of RT-induced genes in Cluster 6 reveals upregulation of TCR-based T-cell activation including bottom: *Gzmb* and *Zap70*. g. Differential gene expression analysis of irradiated versus control macrophage clusters (Cluster 0, 3, 10) followed by Top: GO (MolSigDB) analysis of significantly RT-induced genes reveals induction of interferon signaling and increased M1 macrophage polarization as shown by bottom: violin plots of interferon gene signatures and M1 macrophage gene signatures. h. Differential gene expression analysis of irradiated versus control Cluster 1, Cluster 4, or Cluster MPNST tumor cells following by top: GO analysis (ChEA) of significantly RT-induced genes in Cluster 1 or Cluster 4 reveals enrichment of NFE2L2 targets and bottom: GO analysis (MolSigDB) of significantly RT-induced genes in Cluster 7 shows regulation of immunomodulatory pathways including IL-2/STAT-5 and interferon signaling.

### MPNST clinical outcomes post-RT are affected by tumor purity and CD8+ mediated-immune cell infiltration

While our *in vitro* and *in vivo* analysis supports the importance of IFN signaling and T cell recruitment from the microenvironment as critical for radiation responses in MPNST, the relevance of these putative predictive biomarkers for people with MPNSTs is not clear. We previously identified immunohistochemical patterns associated with radiotherapy response in patients with MPNSTs,^3^ but the predictive significance of specific genetic alterations, methylation signatures, or microenvironmental composition is not well understood. We thus assembled an updated cohort of 30 patients with MPNSTs treated with surgical resection and postoperative radiation who underwent methylation profiling with detailed clinical follow-up. Unsupervised hierarchical clustering identified two epigenetic groups (Figure 4a, Supplementary Figure 7a-b). Epigenetic group 1 tumors had significantly increased copy number variants (CNVs) but no significant difference in age, NF-1 status, or biologic sex (Supplementary Figure 7c-f). Immunomethylomic deconvolution^23,24^ revealed group 1 tumors harbored significantly increased CD4+ T-cells and significantly decreased monocytes, with a trend toward decreased CD8+ T-cells, compared to group 2 tumors, suggesting microenvironmental composition was associated with epigenetic group (Supplementary Figure 7g). Next, we sought to determine whether microenvironmental composition was associated with radiotherapy response as measured by local progression free survival (PFS). Tumor purity estimation using either ABSOLUTE or ESTIMATE methods^25^ demonstrated that low purity tumors with increased microenvironmental signatures were associated with improved local PFS, suggestive of improved radiotherapy response (Figure 4b-c). Similarly, MPNST stratification by CD8+ T-cell infiltration (present versus absent) revealed that patients with tumors harboring CD8+ T-cells were associated with significantly improved local PFS (Figure 4d). Finally, the only recurrent genetic alteration associated with local PFS in this cohort of patients who received radiation was chromosome 9p deletion (Figure 4e). In addition to *CDKN2A/*B, which has been previously implicated in the malignant transformation of MPNSTs, the chromosome 9p21.3 locus contains the interferon gene cluster. Indeed, we observed significantly decreased CD8+ T cell infiltration in 9p lost compared to 9p intact tumors (Figure 4f). Taken together, our human MPNST cohort reinforces the importance of immune microenvironment and T cell recruitment potentially linked to 9p loss in underlying radiation responses in patients with MPNSTs, consistent with our functional genomic and scRNA-seq analysis.

**Figure 4.**
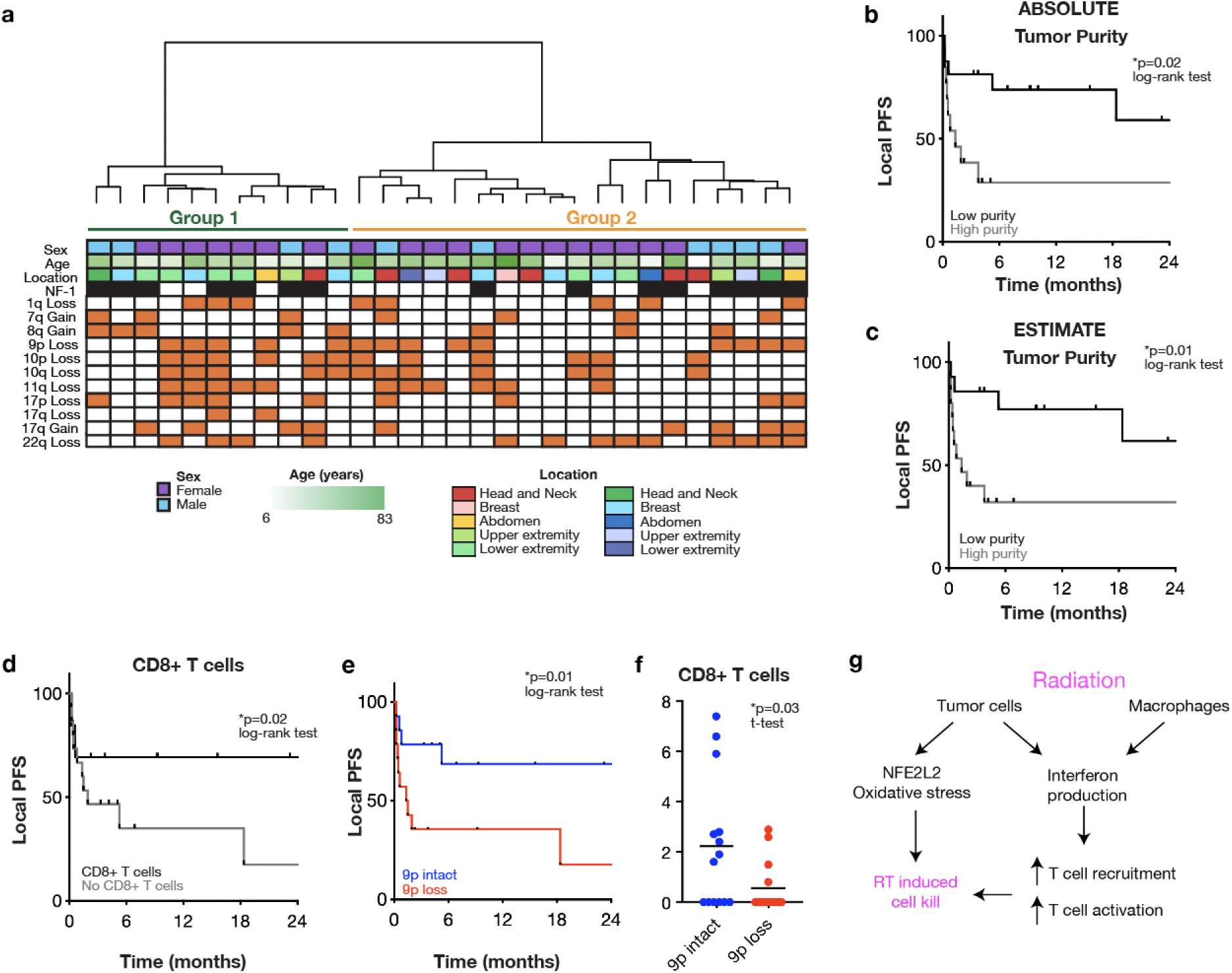
Microenvironmental composition and chromosome 9p loss are predictive of radiotherapy response in a human MPNST cohort. a. Unsupervised hierarchical clustering of DNA methylation arrays from resected human MPNST specimens (n=30) reveals two clusters. Decreased tumor purity reflecting increased microenvironmental composition as measured by b. ABSOLUTE or c. ESTIMATE is associated with improved local progression free survival (PFS) following radiotherapy. d. Methylation cell type deconvolution-based estimate of CD8+ T cells shows the presence of CD8+ T cells is associated with improved local PFS following radiotherapy. e. Chromosome 9p loss is associated with improved local PFS following radiotherapy and f. significantly increased CD8+ T cells. g. Model summarizing tumor cell autonomous and microenvironmental effects of radiation on MPNSTs converging on interferon production underlying T cell recruitment to drive radiation induced cell kill.

### Discussion

MPNSTs are aggressive PNS tumors that are the most common cause of death in people with NF-1. Despite multimodal therapy, clinical outcomes for patients with MPNST remain poor, indicating an unmet need for improved treatment strategies. Here, we combine bulk RNA-sequencing, genome wide CRISPRi screens, and scRNA-seq of immunocompetent mouse MPNST allografts with molecular analysis of human patient MPNST specimens to identify a critical role for interferon signaling and the immune microenvironment in modulating radiation responses (Fig. 4g).

The differential radiation response between NF-1 associated neurofibromas and MPNSTs has important clinical implications given the concern for secondary radiation induced malignancies in people with neurofibromatosis.^5^ Indeed, our data suggest that MPNSTs maintained both a survival and proliferation advantage over pNFs in response to RT and exhibited a distinct anti-apoptotic and anti-inflammatory transcriptional signature across both early and late timepoints compared to pNFs *in vitro*. While the basis for this differential RT response remains to be fully elucidated and validated, one explanation is that *NF1^-/-^* pNFs that lack the additional co-alterations present in MPNSTs, such as chromosome 9p loss or PRC2 mutation, are intrinsically more sensitive to radiotherapy while these same alterations in MPNSTs support a stronger pro-survival response to radiotherapy. However, given the microenvironmental changes observed upon transition from pNF to MPNST,^26,27^ it is also likely that non tumor cell autonomous mechanisms such as IFN signaling underlie both transformation and radiotherapy responses within MPNSTs.

Radiation induced type I IFN signaling, whether from tumor cells or the immune microenvironment, was observed across multiple MPNST models. Prior studies have demonstrated IFN signaling through STAT1 and other IFN stimulated genes is critical for radiotherapy responses.^28–31^ These observations led to clinical trials combining radiation with IFNs that showed some efficacy but significant toxicity,^18^ underscoring the narrow therapeutic window for systemic IFN modulation. One major roadblock to therapeutically leveraging IFNs in cancer has been the complexity of IFN signaling, both with respect to upstream inputs regulating IFN target gene expression and downstream outputs that mediate the effects of IFN activation.^32^ By combining bulk RNA-seq, genome wide CRISPRi screens, and scRNA-seq with a clinically annotated MPNST cohort, our study builds on the connection between IFNs and radiation response by defining specific IFN signaling nodes (*TMEM173, IFNAR1)*, transcription factors (*IRF3, STAT1*), and gene sets required for radiotherapy response in MPNSTs. Indeed, we observed promotion of MPNST radioresistance following knockdown of multiple type I IFN-related genes. From a tumor heterogeneity perspective, we observed type I IFN modulation in both tumor cells and microenvironmental macrophages with associated macrophage polarization toward an M1 state, leading to the recruitment of T-cell populations post-RT *in vivo*. These data underscore the importance of multiple IFN and chemokine sources arising both from the tumor and the microenvironment in determining the anti-tumor immune response. In contrast, our CRISPRi screen data suggests knockdown of type II IFN signaling was noted to promote radiosensitivity, suggesting a potential radioprotective effect of type II IFNs which has been observed in recent work.^33^ Additional validation followed by mechanistic interrogation of why some IFN signaling nodes, but not others, mediate radiotherapy responses will be an important area of future investigation both for diagnostic biomarkers and therapeutic approaches synergizing with radiation.

Analysis of our clinical cohort suggests increased microenvironmental infiltration is associated with improved radiation responses while chromosome 9p loss is associated with decreased radiation response and T-cell infiltration. We also observed differences in the proportion of CD4+ T-cells between methylation clusters and an association between the presence of CD8+ T-cells with radiation response, consistent with previous literature linking radiation induced type I interferons to CD8 T-cell function.^34^ It is important to consider that methylation-based deconvolution is limited in its ability to define distinct immune subpopulations, and orthogonal validation by more robust methods such as flow cytometry, which require tumor samples collected prospectively following radiation, will be an important next step to confirm our methylation-based signatures and better delineate the immune subpopulations that mediate differences in microenvironmental composition. Moreover, given the rarity of MPNSTs and other sarcoma subtypes, observations within our single institution cohort will no doubt require future multi-institutional, ideally prospective, studies.

While chromosome 9p deletion is a well-established event in malignant transformation from pNF to MPNST,^35^ the impact of additional genes lost beyond *CDKN2A/B* within this chromosomal region remains unclear. Notably, the chromosome 9p21.3 region contains a type I IFN gene cluster, and we speculate chromosome 9p loss may thus provide a genetic basis for impairing IFN activation and radiation response in MPNSTs. Indeed, chromosome 9p21 loss is associated with immunotherapy resistance across multiple cancers,^36,37^ and mouse genetic studies further suggest this IFN co-deletion underlies immune evasion.^38^ However, the relationship between 9p loss, with its concomitant IFN co-deletion, and radiotherapy responses remains an area in in need of further investigation. In addition, it will be equally critical to unravel the specific ligand-receptor pairs mediating the crosstalk between MPNST tumor cells and the microenvironment following radiotherapy as well as the specific T-cell subpopulations regulated by these signals.

In sum, the present work suggests multiple mechanisms converge on type I IFN signaling to drive immunomodulation of T-cell populations underlying radiotherapy response in MPNSTs. Future work more precisely defining the mechanisms, regulators, and effectors of type I interferons will be critical to design better tolerated therapies for patients that leverage these immunologic circuits, perhaps by directly targeting STING.^39^ In addition, given the limited number of available MPNST samples from patients who had received adjuvant radiotherapy, our clinical analysis is likely underpowered, and further multi-institutional efforts will be important to determine if these observations hold up in larger cohorts. Finally, our data establish the preclinical rationale for testing combination approaches with radiotherapy designed to stimulate interferon secretion, which can perhaps be best achieved with intralesional approaches^40^ such as through intra-tumoral oncolytic virus administration,^41^ as a rational approach to improve radiation responses for these aggressive tumors.

## Acknowledgments

We thank Christine Pratilas from Johns Hopkins University for JH-2-002 MPNST cells and Angie Hirbe from the Washington University in St. Louis for JW18.2 and JW23.3 *Nf1/Tp53* mouse MPNST cells. We also thank Joanna Phillips, Anny Shai and the staff of the UCSF Brain Tumor Center Biospecimen and Pathology Core for histological staining and Eric Chow and the staff of the UCSF Center for Advanced Technology for sequencing. We thank the members of the Vasudevan Laboratory, Kim Rickman, Alex Perez, and Mary Helen Barcellos-Hoff for critical comments on the manuscript, and we thank Will C. Chen for assistance with mouse irradiation experimental setup. H.N.V. was supported by a Francis Collins Scholar Fellowship from the Neurofibromatosis Therapeutic Acceleration Program (NTAP).

## Author contributions statement

All authors made substantial contributions to the conception or design of the study; the acquisition, analysis, or interpretation of data; or drafting or revising the manuscript. All authors approved the manuscript. All authors agree to be personally accountable for individual contributions and to ensure that questions related to the accuracy or integrity of any part of the work are appropriately investigated, resolved, and the resolution documented in the literature.

I.Z. designed, performed, and analyzed all experiments and bioinformatic analyses. J.C., K.M., and J.P. performed QPCR, cell culture experiments including colony forming assays, and mouse MPNST allograft experiments. K.M. assisted with assembling human tumor resection specimens, bioinformatic analysis, and pathologic review. S.P. performed single-cell RNA sequencing and assisted with bioinformatic analysis. V.M., S.E.B., and L.J. provided key insight into study design and provided clinical data. M.P. helped assemble tumor resection specimens, provided clinical data, performed pathologic review, and assisted with study design. J.L. provided critical experimental oversight including in design of CRISPR screens and assisted with bioinformatic analysis. H.N.V. conceived, designed, and supervised the study.

## Competing interests statement

The authors declare no competing interests.

## Methods

This study complied with all relevant ethical regulations and was approved by the UCSF Institutional Review Board (13-12587, 17-22324, 17-23196, 18-24633, 22-37134). As part of routine clinical practice at UCSF, all patients included in this study signed an informed waiver of consent to contribute de-identified data to scientific research projects.

### Tissue culture

Patient-derived neurofibroma (NF95.11b) or human MPNST (JH002-2, ST88-14) cell lines were obtained from the Neurofibromatosis Therapeutic Acceleration Program or American Type Culture Collection. Mouse MPNST cell lines (JW18.2, JW23.3) were a gift from Dr. Angie Hirbe. Cell lines were grown in Dulbecco’s Modified Eagle Medium (#11960069, Life Technologies) with 10% FBS and 1X Pen-Strep (#15140122, Life Technologies). Cell lines were regularly tested and verified to be mycoplasma negative (#LT07-218, Lonza).

### Flow cytometry

For proliferation assays, cells were stained with Cell Trace Yellow (Thermo Fisher) according to manufacturer’s protocol. Cells were then treated with a single-fraction of radiation using a X-Rad 320 irradiator (Precision X-ray) or no radiation and incubated for 96 hours after completion of radiation. For survival assays, cells were irradiated with a course of daily 2 Gy fractions and stained with DRAQ7 (Fisher Scientific) 96 hours after irradiation. Flow cytometry and FACS were performed on a Becton Dickinson (BD) FACS Aria Fusion and analyzed using FlowJo software. Samples of cell suspension from each tumor sample were stained with PE-conjugated anti-murine CD45 (need ID#), PECy7-conjugated anti-murine CD3e (17A2, BD Biosciences), BUV-conjugated anti-murine CD4 (GK1.5, BD Biosciences), and BV786-conjugated anti-murine PD-1 (29F.1A12, BD Biosciences). Flow cytometry was performed as above.

### Nucleic acid extraction and qRT-PCR

RNA was extracted from cell lines using the RNeasy Mini Kit (#74106, QIAGEN) according to manufacturer’s instructions, and cDNA was synthesized using the iScript cDNA Synthesis kit (#1708891, BioRad). Real-time QPCR was performed using PowerUp SYBR Green Master Mix (#A25918, Thermo Fisher Scientific) on a QuantStudio 6 Flex Real Time PCR system (Life Technologies) and analyzed using the double delta method as previously reported. The following QPCR primers were used: GAPDH-F (5′-GTCTCCTCTGACTTCAACAGCG-3′), GAPDH-R (5′-ACCACCCTGTTGCTGTAGCCAA-3′), KEAP1-F (5’-CGGGGACGCAGTGATGTATG-3’), KEAP1-R (5’-TGTGTAGCTGAAGGTTCGGTTA-3’), IRF3-F (5’-GAGAGCCGAACGAGGTTCAG-3’), IRF3-R (5’-CTTCCAGGTTGACACGTCCG-3’), TMEM173-F (5’-CCAGAGCACACTCTCCGGTA-3’), TMEM173-R (5’-CGCATTTGGGAGGGAGTAGTA-3’), IFNAR1-F (5’-AACAGGAGCGATGAGTCTGTC-3’), IFNAR1-R (5’-TGCGAAATGGTGTAAATGAGTCA-3’), STAT1-F (5’-CAGCTTGACTCAAAATTCCTGGA-3’), and STAT1-R (5’-TGAAGATTACGCTTGCTTTTCCT-3’).

### Bulk RNA-sequencing

Library preparation was performed using the TruSeq RNA Library Prep Kit v2 (#RS-122-2001, Illumina). 50 bp single end reads were sequenced on an Illumina HiSeq 2500 or NovaSeq to a minimum depth of 25M reads per sample at Medgenome, Inc. Following quality control of FASTQ files with FASTQC, trimming of adapter sequences, reads were filtered to remove bases that did not have an average quality score of 20 within a sliding window across 4 bases (http://www.bioinformatics.babraham.ac.uk/projects/fastqc/) was performed prior to mapping. Reads were mapped to the appropriate reference genome (hg19) using HISAT2 with default parameters.^42^ Transcript abundance estimation in transcripts per million (TPM) and differential expression analysis were performed using DESeq2.^43^ Principal component analysis was performed in R using the prcomp function. Differentially expressed transcripts with an adjusted p-value<0.1 were identified and filtered based on an expression cutoff (TPM>1) and a fold change threshold (log2FC>1) to prioritize biologically relevant gene sets.

### CRISPRi cell line generation and genome-wide screening

Lentivirus containing UCOE-SFFV-dCas9-BFP-ZIM3 (#188777, Addgene) was produced from transfected HEK293T cells with packaging vectors (pMD2.G #12259, Addgene, and pCMV-dR8.91, Trono Lab) following the manufacturers protocol (#MIR6605, Mirus). JH-2-002 cells were stably transduced to generate parental JH-2-002^dCas9-Zim3-BFP^ cells and selected by flow cytometry using a SH800 sorter (Sony). Subsequent gene specific knockdowns were achieved by individually cloning single-guide RNA (sgRNA) protospacer sequences into the pCRISPRia-v2 vector (#84832, Addgene) between BstXI and BlpI restriction sites. All constructs were validated by Sanger sequencing of the protospacer region. The following protospacers were used: sgNTC (GTGCACCCGGCTAGGACCGG), sgKEAP1-1 (GGCCCTGGCCTCAGGCGGTA), sgKEAP1-2 (GTGGAGCCGAGGCCCCCCGA), sgIRF3-1 (GGGAGGCGTCTACACTGAGG), sgIRF3-2 (GGGGTGGACTCCGTAGATGG), sgTMEM173-1 (GAGAGCAGCCAGTGTCCGGG), sgTMEM173-2 (GGGTGCCCAGCCACTCCCAG), sgIFNAR1-1 (GTAACTGGTGGGATCTGCGG), sgIFNAR1-2 (GATGTAACTGGTGGGATCTG), sgSTAT1-1 (GGCAGGAAAGCGAAACTACC), and sgSTAT1-2 (GCTGCGCAGAGTCTGCGGAG). Lentivirus was generated as described above and cells were selected to purity using 1-5 μg/mL puromycin for at least 5 days.

For genome wide CRISPRi screening, we used a compact and highly active barcoded sgRNA library that was optimized through aggregation of 126 genome wide CRISPRi screens, established sgRNAs targeting essential genes, and machine learning prediction algorithms.^24^ This genome-wide dual sgRNA library has been previously validated through multiple growth-based screens as well as through confirmation of on-target gene repression using perturb-seq, exhibiting 82-92% median target knockdown^24^. This genome-wide dual sgRNA library containing the top 2 on-target sgRNAs for 23,483 genes was cloned into the library expression vector pU6-sgRNA Ef1alpha Puro-T2A-GFP derived from pJR85 (#140095, Addgene) and modified to express a second sgRNA using the human U6 promoter as previously described.^24,59^ Knockdown efficiency of all guide sequences in this genome-wide sgRNA library was previously validated in K562 cells as part of a genome wide Perturb-seq database, and this data is publicly available at https://gwps.wi.mit.edu/. 1137 non-targeting sgRNA pairs were also included as negative controls in the screen. To generate lentiviral pools, HEK293T cells were transfected with the sgRNA library along with packaging plasmids as described above, and viral supernatant was collected 72 hours following transfection. Lentiviral libraries were transduced into JH-2-002^dCas9-KRAB-BFP^ cells, cultured for 2 days following infection, selected in 1 μg/mL puromycin for 2 days, and then allowed to recover in 10% FBS in DMEM for 1 day. Cells were then sorted for GFP expression using FACS to obtain a pure population of 1x10^7^ cells, and cells were subsequently cultured to allow for 1x10^7^ cells per biological replicate. Two pellets of 1x10^7^ cells were subsequently frozen down at this “T0” timepoint. The screen was subsequently carried out in biologic triplicate with two experimental groups, one receiving 2 daily doses of 2 Gy and one receiving no radiation. Genomic DNA from all cells was harvested at the T14 (14 days) endpoint and sgRNA barcodes were amplified and processed for sgRNA abundance library preparation using Q5 High-Fidelity DNA Polymerase (NEB) and sequenced on an Illumina NovaSeq 6000 as previously described^59^.

Enrichment or depletion of sgRNA abundances were determined by down sampling trimmed sequencing reads to equivalent amounts across all samples, and then calculating the log2 ratio of sgRNA abundance in experimental conditions to sgRNA abundance in control conditions at T14, or between sequencing reads from T14 and T0 timepoints within experimental or control conditions. Specifically, we computed normalized log2 ratios for radiation treated sgRNA abundance at T14 as compared to control T14 abundances in order to identify mediators of radiation responses and computed normalized log2 ratios for untreated sgRNA abundance at T14 compared to T0 to identify regulators of cell fitness independent of treatment. Any sgRNA’s not represented with at least 50 normalized sequencing reads across all replicates were excluded from analysis.^24^ Statistical significance was calculated using the Wald test comparing replicates across conditions without a log2 fold change threshold. The screen was analyzed to identify significantly enriched or depleted guides with either vehicle treatment or radiation with the latter being the focus for genetic mediators of radiation response. Hits were prioritized by normalizing log2 ratios to the total number of population doublings in the screen. These phenotype log2 ratios were used for subsequent analysis and visualization. Genes were filtered at an adjusted p-value<0.05 for statistical significance.

### Mouse subcutaneous allograft tumors

The study was approved by the UCSF Institutional Animal Care and Use Committee (AN200569-00) and all experiments were conducted in compliance with institutional and governmental regulations. Subcutaneous allografts were performed by implanting 5 million JW18.2 or JW23.3 MPNST allografts cells into the flanks of 5–6-week-old female C57/B6 wildtype mice (Charles River). For radiation treatments, the desired radiation was delivered using the XRAD 320 irradiator, and custom 3D printed lead shielding was used to prevent radiation delivery to regions outside the palpable, visualized subcutaneous tumor. Tumors were measured using calipers 3 times per week.

### DNA methylation profiling

Methylation profiling was performed using the Illumina EPIC array platform. Archival FFPE tissue blocks were screened, and DNA was extracted using the Qiagen FFPEasy kit and 1500 ng DNA was loaded per sample. DNA concentrations down to a minimum of 20ng/µl (total: 1.5µg) were considered acceptable for methylation array profiling. DNA quality was assessed by spectrophotometry, and clean-up was performed as needed using DNA precipitation. Preprocessing, normalization, and downstream analysis were performed in R using the minfi Bioconductor package, as previously reported.^21,44–46^ Data were normalized via functional normalization and filtered based on the following criteria: (i) removal of probes targeting the X and Y chromosomes, (ii) removal of probes containing a common single nucleotide polymorphism (SNP) within the targeted CpG site or on an adjacent basepair, and (iii) removal of probes not mapping uniquely to the human reference genome. Unsupervised hierarchical clustering (Euclidean distance, complete linkage method) was performed using the top 2,000 most variable probes as previously described.^47^ β values were used for visualization of methylation levels (β=methylated/[methylated+unmethylated]) and M values were used for statistical analysis (M=log2[methylated/unmethylated]). ConsensusClusterPlus (v 1.62.0) analysis (hierarchical clustering, k range 2 to 10, 1000 repetitions) was used to assess optimal cluster size and stability.

### Single cell RNA-sequencing

For mouse allograft single-cell RNA sequencing, tumors were minced with sterile razor blades and then enzymatically dissociated with papain (#LS003, Worthington) at 37 degrees Celsius for 45 minutes. Samples were subsequently centrifuged for 5 minutes at 500g, resuspended in RBC lysis buffer (#00-4300-54, eBioscience), incubated for 10 minutes at room temperature, and then resuspended in 5% FBS in PBS. Cell suspensions were serially filtered through 70 μm and 40 μm filters before being resuspended again in 5% FBS in PBS for manual cell counting using a hemacytometer. A total of 10,000 cells were loaded per single-cell RNA sequencing sample onto the 10x platform using the Chromium Single Cell 3’ Library & Gel Bead Kit v3.1 on a 10X Chromium controller (10X Genomics) per the manufacturer recommended default protocol and settings. Samples were sequenced on an Illumina NovaSeq at the UCSF Center for Advanced Technology, and the demultiplexed FASTQ files were processed using CellRanger for alignment to the mm10 reference genome, identification of empty droplets, and determination of a count threshold. Cellranger generated filtered feature matrices were imported into a Seurat object. All downstream analyses were performed with Seurat v4.4.^48^ For quality control, data were filtered on a per sample basis to remove outliers in gene count, UMI count, mitochondrial genes, and ribosomal genes. The individual count matrices were normalized by SCTransform v2. Scanorama (https://github.com/brianhie/scanorama)^49^ was used to perform data integration across datasets and cluster number optimization was performed by comparing multiple cluster resolutions using Clustree and the selected cluster resolution was examined by silhouette-width analysis, which reported a mean width per cluster larger than 0. Genes differentially expressed in each cluster were identified using FindAllMarkers function (cutoff: min.pct = 0.25, logfc.threshold = 0.25, min.diff.pct = 0.1). Tumor vs non-tumor cell microenvironment designation was based on *Xist* gene expression, which was expressed in female host mouse cells but not male MPNST tumor cells, scType automated cell identification,^22^ and manual inspection of differentially expressed gene markers as previously described.^50^

### Statistical Analysis

All experiments were performed as repeated, independent biologic replicates, and statistics were derived from biologic replicates. The number of biologic replicates is indicated in each panel or figure legend. No statistical methods were used to predetermine sample sizes. Considering the rarity of MPNSTs and accounting for the number of genomic approaches used in this study, our total cohort size is similar to prior publications^7–9^. The clinical samples used were retrospective and non-randomized, and all samples were equally interrogated within the constraints of sufficient tissue for each analytical method. Cells and animals were randomized to experimental conditions, and no clinical, molecular, cellular, or animal data points were excluded from analysis. Unless otherwise specified, data are plotted as mean with error bars that represent the standard error of the mean. The statistical tests of choice were selected based on the input data and are noted in the methods and figure legends. All statistical tests were two-sided. Where appropriate, multiple hypothesis testing corrections were performed. Statistical significance thresholds are indicated in each figure legend and exact p-values are provided whenever possible.

## Data Availability Statement

Human tumor DNA methylation, bulk RNA-sequencing, and single cell RNA sequencing reported in this manuscript have been deposited in the NCBI Gene Expression Omnibus under GEO records GSE290457, GSE290674, GSE289110 and GSE289111.

## Code availability

The open-source software, tools, and packages used for data analysis in this study, as well as the version of each program, were ImageJ (v2.1.0), R (v3.5.3 and v3.6.1), cellranger (v6.1.2), Seurat R package (v4.4.0), Clustree (v0.5.0), Scanorama (v1.7.3), minfi (Bioconductor v3.10), ConsensusClusterPlus (Bioconductor v3.10), Heatmap.2 R package (gplots v3.13), and ggplot2 (v3.4.3). No custom software, tools, or packages were used. CRISPRi screen analysis code is available at https://github.com/liujohn/CRISPRi-dual-sgRNA-screens/blob/main/module2/PhenotypeScores.R.

## Supplementary Figures

**Supplementary Figure 1.**
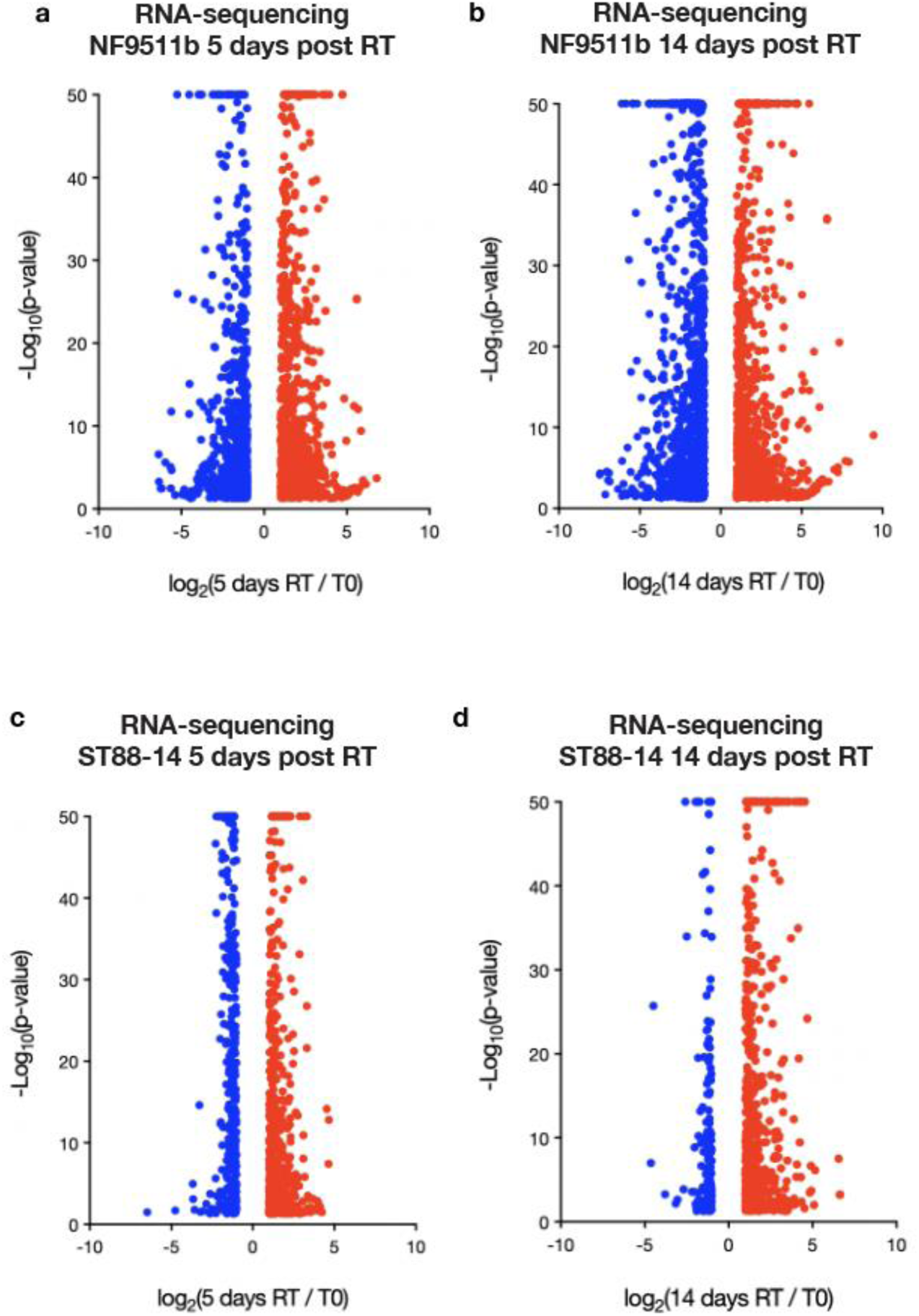
Bulk RNA-sequencing analysis of NF9511b pNF or ST88-14 MPNST cells at early (5 days) or late (14 days) timepoints following radiation. Volcano plots depicting significantly modulated genes in NF9511b pNF cells at a. 5 days (793 upregulated genes, 757 downregulated genes) or b. 14 days (937 upregulated genes, 1178 downregulated genes) following radiation and in ST88-14 MPNST cells at c. 5 days (651 upregulated genes, 652 downregulated genes) or d. 14 days (605 upregulated genes, 186 downregulated genes) following radiation. Genes were considered significant at p < 0.05 and - log10(p-value) scores were capped at 50 for visualization purposes.

**Supplementary Figure 2.**
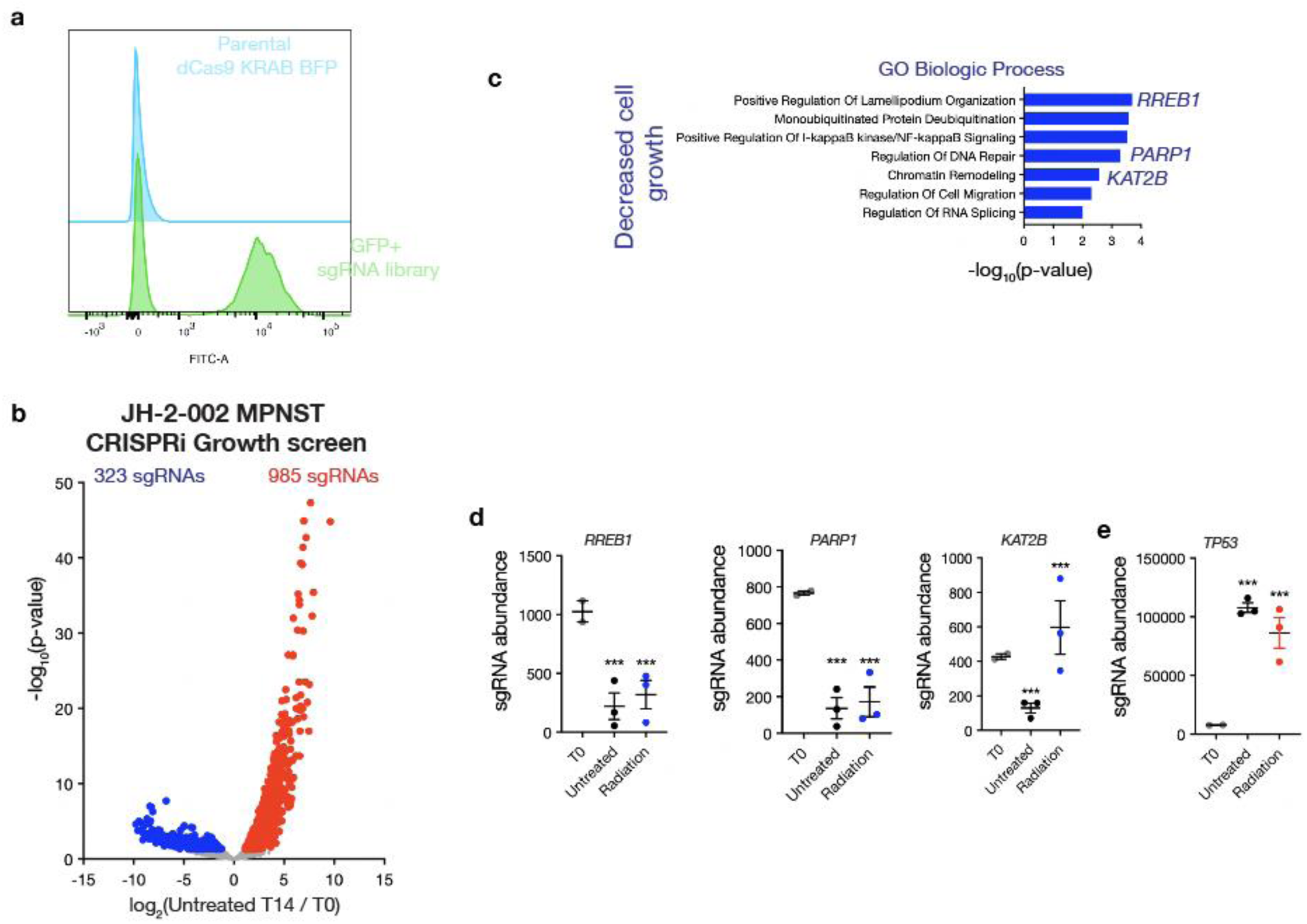
Genome wide CRISPRi growth screen (T14 untreated compared to T0) analysis in JH-2-002 MPNST cells. a. Flow cytometry demonstrating gating parameters for GFP+ sgRNA library (bottom) compared to negative control, un-transduced dCas9 KRAB BFP parental cell line (top). b. Volcano plot depicting 323 significantly depleted sgRNAs (blue) and 985 significantly enriched sgRNAs (red) mediating MPNST cell growth at T14 compared to initial T0 timepoint. c. Gene ontology analysis of genes required for MPNST cell growth reveals genes and processes known to be important in MPNSTs including cytoskeletal Ras target genes (*RREB1*), DNA repair (*PARP1*), or chromatin remodeling (*KAT2B*). e. In addition, sg*TP53* was amongst the top hits mediating increased MPNST cell growth, serving as a positive control. *** p<0.01 compared to T0.

**Supplementary Figure 3.**
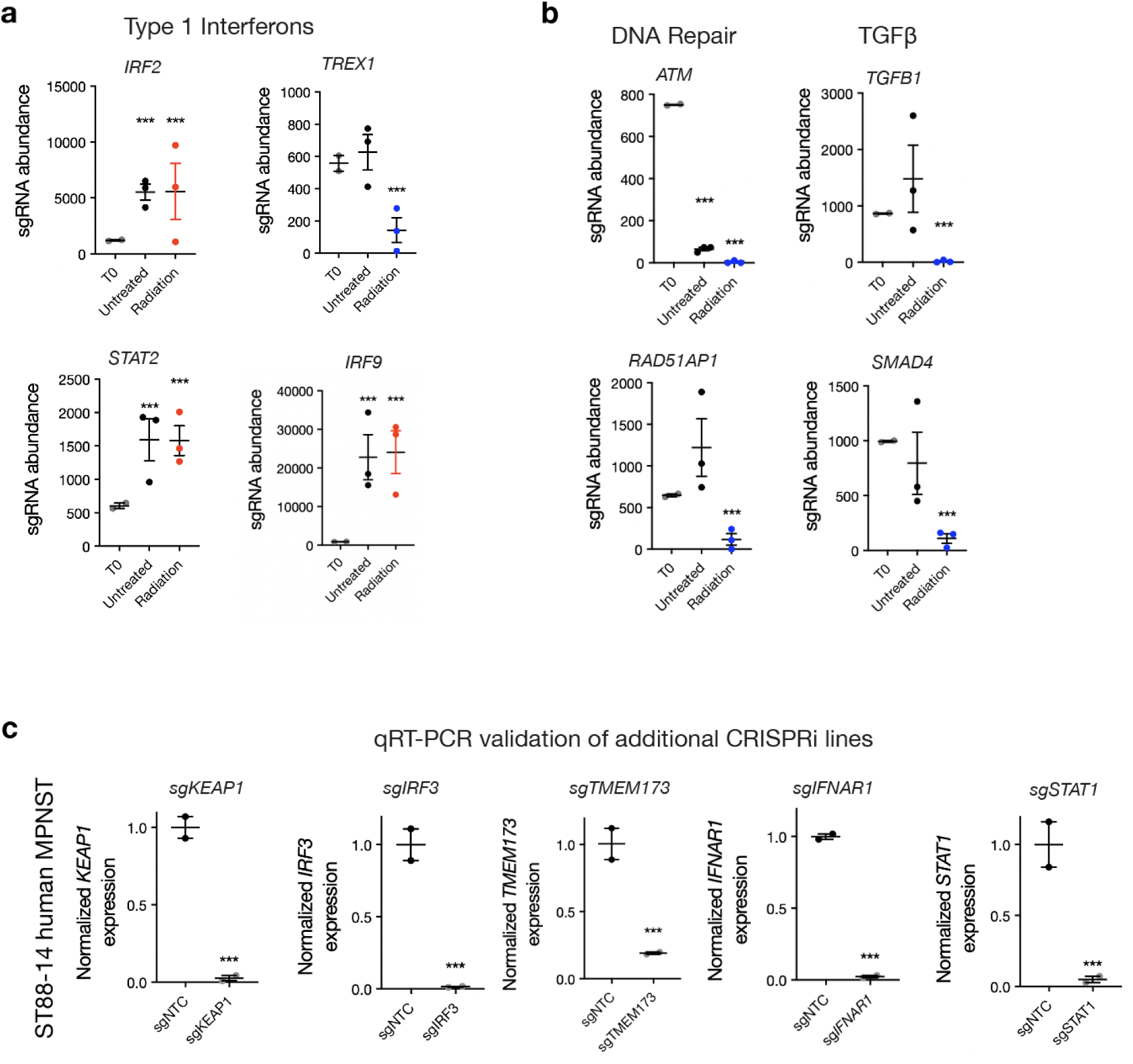
Selected CRISPRi screen hits significantly modulating radiation response. a. Multiple components of type 1 interferon signaling (*IRF2, STAT2, IRF9)* mediate radiation resistance while repression of the negative regulator of STING *TREX1* significantly mediates radiation sensitivity. b. Consistent with their well-established roles in cell autonomous radiation response, DNA repair (*ATM, RAD51AP1)* and TGFβ signaling (*TGFB1, SMAD4*) mediate radiosensitivity. c. qRT-PCR validation in ST88-14 MPNST cells of selected JH-2-002 CRISPRi screen hits (*sgKEAP1, sgIRF3, sgTMEM173, sgSTAT1)*.

**Supplementary Figure 4.**
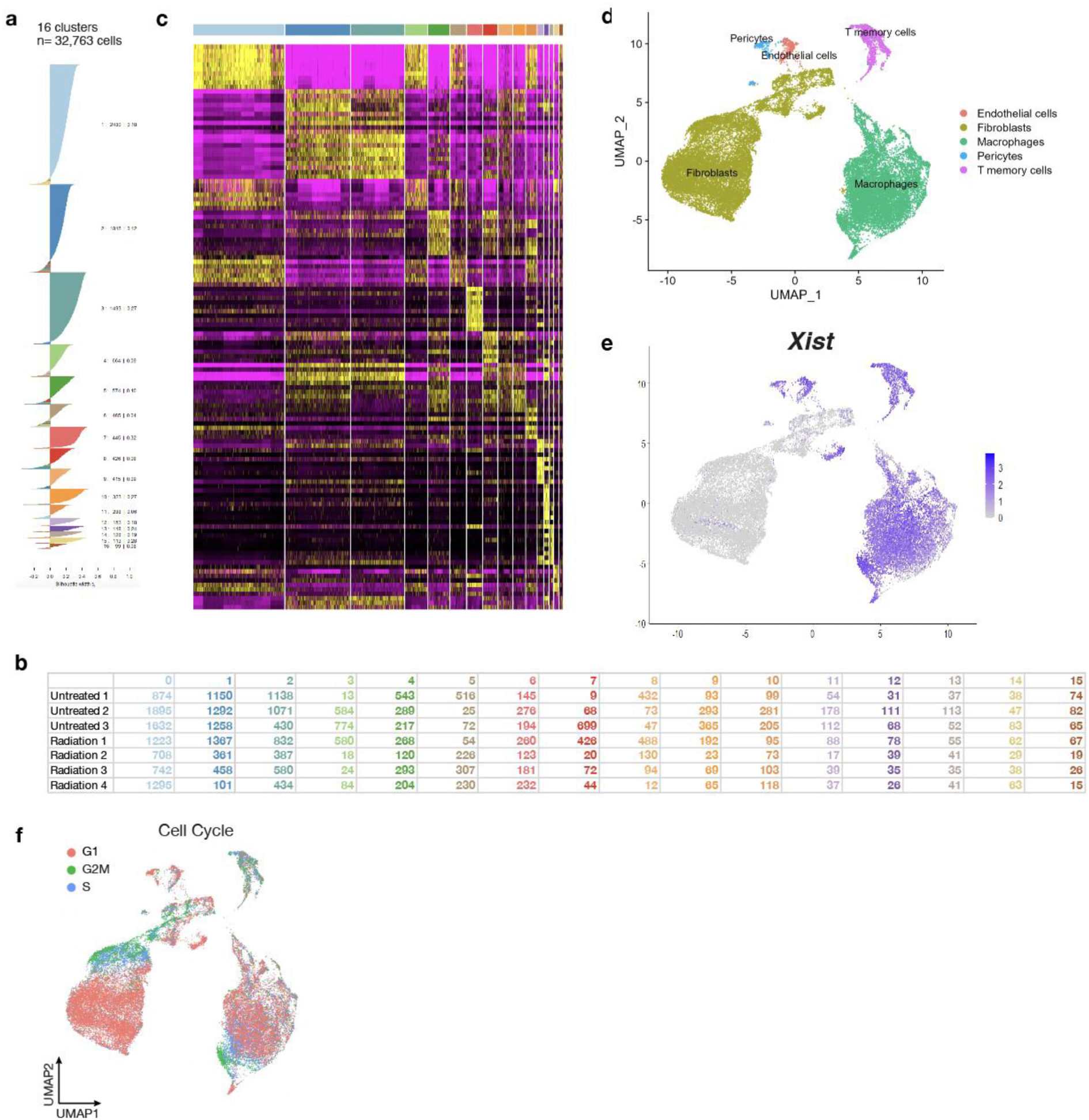
Single cell RNA-sequencing (scRNA-seq) of irradiated and control mouse MPNST allograft tumors implanted subcutaneously in C57/B6 mice. a. Silhouette analysis reveals 16 clusters across 32,763 cells across b. 3 untreated and 4 irradiated tumors. c. Heatmap for cluster marker genes across all clusters integrated with d. automated cell type assignment (SCtype), e. *Xist* expression to define female host mouse microenvironment cells, and f. cell cycle expression signature leads to robust cell type classification.

**Supplementary Figure 5.**
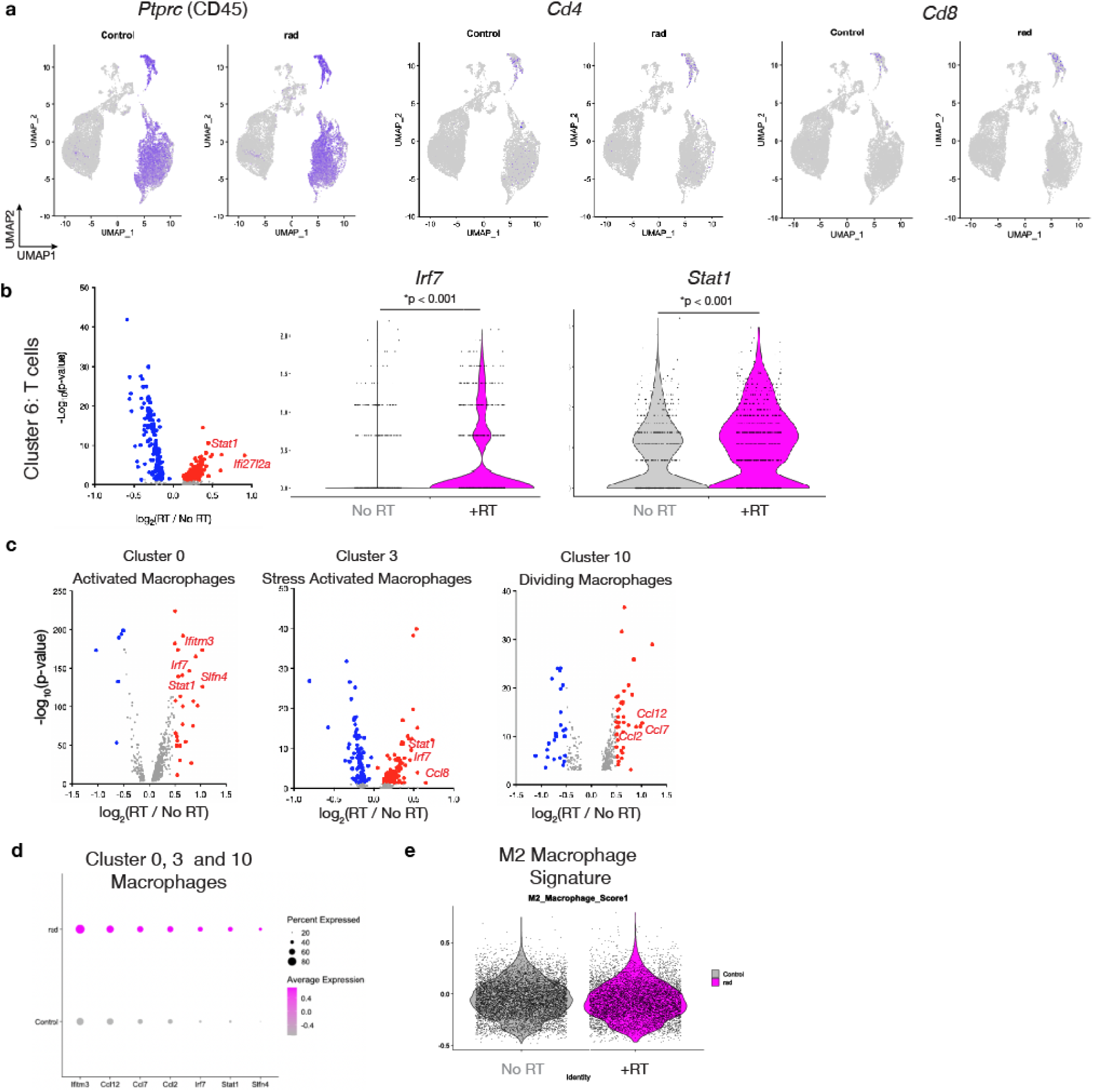
Differential expression analysis of non-tumor T/NK cells (Cluster 6) and macrophages (Clusters 0, 3, and 10). a. *Ptprc* (CD45) expression as a pan leukocyte marker identifies all cells of the leukocyte lineage while CD4/CD8 is restricted to Cluster 6. b. Differential gene expression analysis between irradiated (RT) and control (no RT) Cluster 6 T/NK cells reveals induction of T cell activation and interferon gene circuits (*Irf7, Stat1*) following radiation. c,d. Differential gene expression analysis between irradiation (RT) and control (no RT) macrophage clusters reveals induction of interferon gene circuits (*Irf7, Ifitm3, Stat1)* and multiple chemokines (*Ccl2, Ccl7, Ccl8, Ccl12*) following radiation. e. Irradiation does not significantly alter M2 macrophage polarization.

**Supplementary Figure 6.**
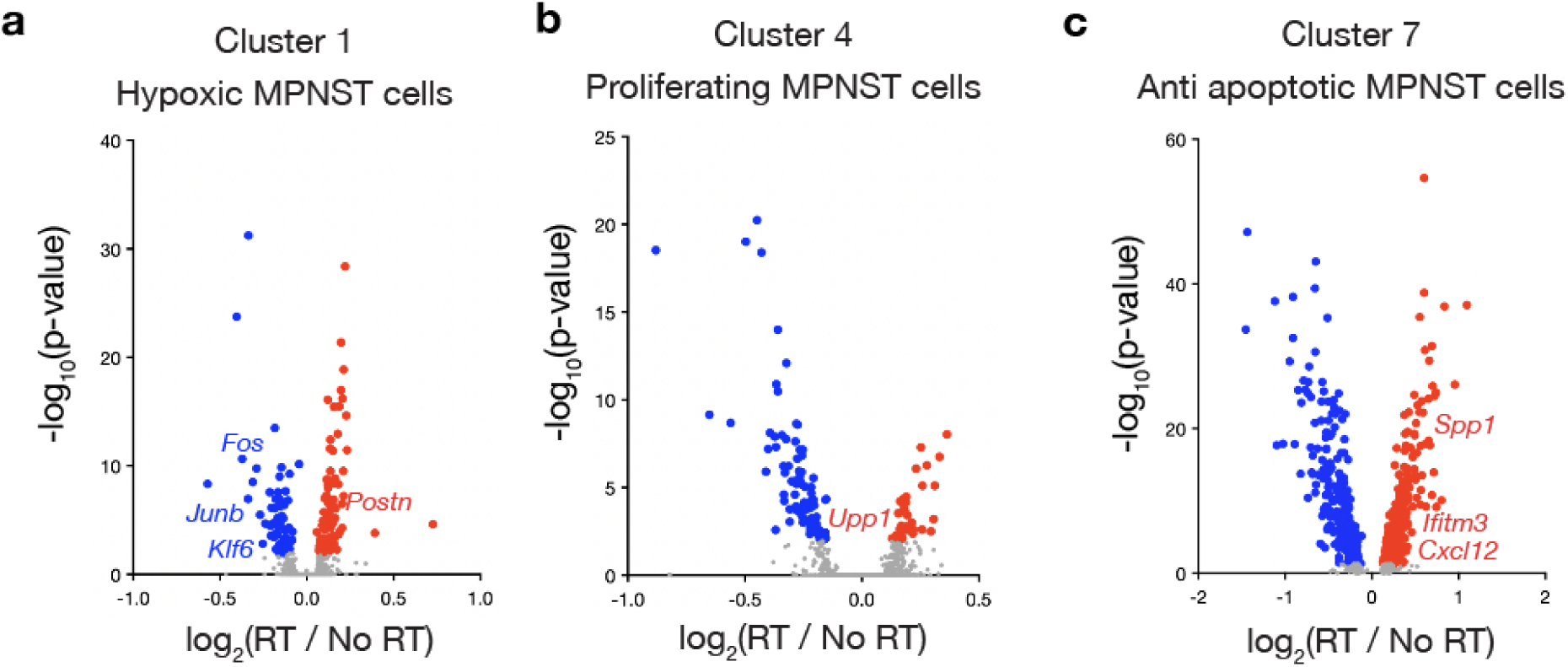
Differential expression analysis of a. hypoxic MPNST cells (Cluster 1), b. proliferating MPNST cells (Cluster 4), or c. anti-apoptotic MPNST cells (Cluster 7) reveals induction of interferon signaling *(Spp1, Ifitm3*) and chemokine secretion (*Cxcl12*) in Cluster 7 cells following irradiation.

**Supplementary Figure 7.**
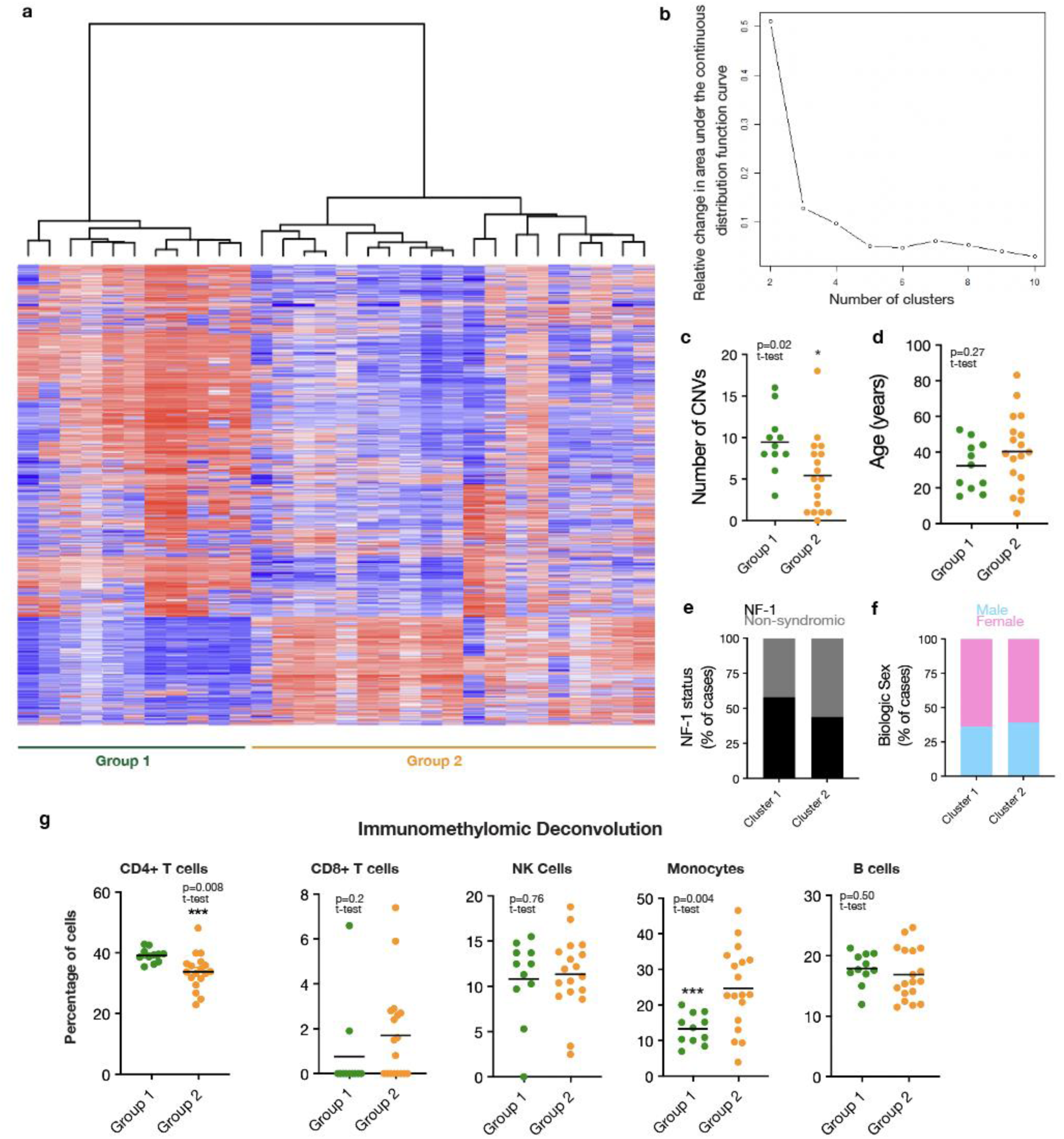
Methylation array analysis of human patient MPNST resection specimens treated with postoperative radiation therapy. a. Unsupervised hierarchical clustering heatmap of top 2,000 most variable probes identifies b. two epigenetic groups based on the change in area under the curve for consensus cumulative distribution function (CDF), which shows minimal appreciable increase for greater than 2 clusters. c. Group 2 tumors have significantly increased CNVs but no significant difference in d. age, e. NF-1 status, or f. biologic sex. g. Immunomethylomic deconvolution of different microenvironmental cell types revealed Group 1 tumors had significantly increased CD4+ T-cells, significantly decreased monocytes, and a trend toward decreased CD8+ T-cells.

## Supplementary Tables

**Supplementary Table 1.** Normalized gene expression values for bulk RNA-sequencing of irradiated plexiform neurofibroma NF9511b or MPNST ST88-14 cells.

**Supplementary Table 2.** Analysis of MPNST JH002-2 CRISPRi screen comparing sgRNA enrichment between cells treated with the radiation (2 Gy x 2 fractions) or no treatment for 10 days

**Supplementary Table 3.** Cluster marker gene list for scRNA-sequencing of tumor cells from irradiated 2 Gy x 5 fractions (n=4) or untreated (n=3) JW23.3 MPNST subcutaneous allografts.

## Notes

### Competing Interest Statement

The authors have declared no competing interest.

